# Mental health explains individual deviations from normative range in cognition-associated brain states

**DOI:** 10.1101/2021.03.12.435017

**Authors:** Wei Zhang, Diego Vidaurre, Janine Diane Bijsterbosch

## Abstract

Links between cognitive deficits and psychiatric disorders have been studied predominantly at the group level, leaving unique individual characteristics largely elusive. Here, we applied normative modeling to UK Biobank data (N=18,634) and estimated the individual-level interplay of large-scale brain networks over time (i.e., dynamic brain state) as a function of general cognitive ability. Abnormality in such brain states was linked to individual variation in mental health. Specifically, brain state measures including fractional occupancy that indicates the brain state probability over time were estimated using a Hidden Markov Model, followed by a Gaussian process regression to estimate the normative range of these brain state measures from general cognitive ability. Abnormality scores per participant were quantified to represent the degree of deviations relative to the estimated population norm. We found significant associations between the abnormality scores of several brain states and individuals’ overall mental health. Our findings suggest potential impact of mental health on dynamic brain states that subserve cognitive functions and shed light on the relevant brain mechanisms underlying cognitive deficits in mental illness.

## Introduction

Cognitive functions are a set of mental abilities such as problem solving and decision making that are required to perform daily tasks. The ability to execute such functions has been shown to vary across the lifespan (*1*), and from one condition to the other (*2*). In fact, impairment in such ability (i.e., cognitive deficit) is recognized as a common symptom present in the majority of psychiatric disorders, including schizophrenia (*3*), depression (*4*) and bipolar disorder (*5*). The observed behavioral abnormalities in those disorders are often attributed to various cognitive functions including learning, attention, emotion regulation and cognitive control (*6*). Aberrant functional connectivity patterns, in addition to other neurobiological measures, have been linked to cognitive deficits in psychiatric disease groups (*7*). A recent meta-analysis study revealed that altered functional connectivity (FC) in large-scale neurocognitive networks were common across psychiatric diagnoses, suggesting a shared mechanism of brain network interactions underlying abnormalities in generalized cognitive ability (*8*). These findings point to the FC of large-scale networks as the mechanistic basis of cognitive ability and suggest that the relationship between FC and cognition may vary as a function of mental health.

Interestingly, recent studies on FC suggest that in contrast to traditional static functional connectivity, patterns of large-scale brain networks over time that capture time-varying functional connectivity might be of greater relevance to studying individual differences in cognitive functions (*9*–*11*). Unlike static FC, which averages neural fluctuations over time, dynamic connectivity estimates temporal fluctuations in functional connectivity and provides additional information about temporal reconfiguration of functional resources (*12*). Growing evidence suggests that dynamic changes in the resting-state FC are involved in various cognitive processes, including mind-wandering, fluctuations in arousal, vigilance, and perceptual performance (*13, 14*). Recent studies linking temporal dynamics of large-scale brain networks to cognitive and behavioral traits further revealed that such network temporal properties often had better predictions on cognitive traits than static FC (*9*), explain unique aspects of cognition (*11*), and that dynamic connectivity can outperform static FC in terms of capturing trial-wise behavioral variability in cognitive tests (*10*). These findings highlight the interactions of brain networks over time as a potentially more sensitive marker than the static FC to capture inter-individual differences in cognition.

Brain network dynamics have also been linked to various psychiatric and neurological disorders. However, almost all reported studies so far have focused on group-level comparisons between patients and healthy controls (*15*). Similarly, research into the role of mental illness in cognitive functions is also predominantly based on case versus control comparisons or clinical subgroup contrasts (*16*). These studies therefore only provide group-level inference that essentially overlook unique individual characteristics. To obtain more representative measures of each individual and to eventually identify meaningful biomarkers, we therefore need more personalized approaches (*16*–*18*).

In this study, we adopt individualized methods to investigate the interactions of large-scale brain networks over time (i.e., brain network state with temporal information) that are associated with cognitive ability, and their relationships to mental health. Specifically, we leverage normative modeling to estimate individual-specific brain state measures from cognitive ability. Analogous to normative growth charts used in pediatric medicine, normative models locate each individual on the normative range in terms of the estimated variables (e.g., brain state measures) and provide statistical inference at the individual subject level (*19*). Deviations from the estimated normative range per individual (i.e., abnormality score) can be then quantified and linked to relevant variables (e.g., individual’s mental health). Normative modeling has been increasingly applied to study associations between brain functions and behavior, particularly for clinically relevant conditions where significant heterogeneity may exist that cannot be captured by categorical partitioning of the cohort into traditional patient-control groups (*20*–*23*). Here, we apply normative modeling to a population cohort to investigate whether individual-level abnormality in brain state measures as estimated from cognitive ability can be attributed to overall mental health.

## Results

### General cognitive ability and mental health state

To summarize general cognitive ability and overall mental health, we performed principal component analyses across a set of cognitive tests and mental health self-report questions. For cognitive tasks, the first principal component derived from all items explained 33.75% of variance, and the first principal component derived from mental health questionnaire items explained 24.08% of variance. Individual scores for these first components per participant were used to indicate general cognitive ability and overall mental health, respectively. This approach has been widely applied to approximate general intelligence (i.e., g-factor) and generalized dimensional symptom levels across psychiatric disorders (i.e., p-factor). The loadings (i.e., eigenvalues) of each individual item on the principal components for cognitive ability and mental health are shown in *Figure S1*.

### Brain state measures

Hidden Markov Modeling (HMM) was performed at the group-level using the concatenated data of all participants’ individual timeseries from 55 components, as defined with independent component analysis, to estimate the interactive patterns of brain networks over time (i.e., brain states). In accordance with previous studies, twelve brain states were inferred from the model, each of which is represented by a multivariate Gaussian distribution, as described by the mean and covariance. Different brain networks, including the default mode and sensorimotor networks were engaged in these inferred brain states (*Figure 1*; also see full brain activation maps in *Figure S2*). Fractional occupancy (FO) of each inferred brain state and the switching rate (SR) across twelve brain states were calculated for each participant to indicate the time they spent in each individual brain state and the overall stability of these brain states across time, respectively (*Figure 2*).

**Figure 1.**
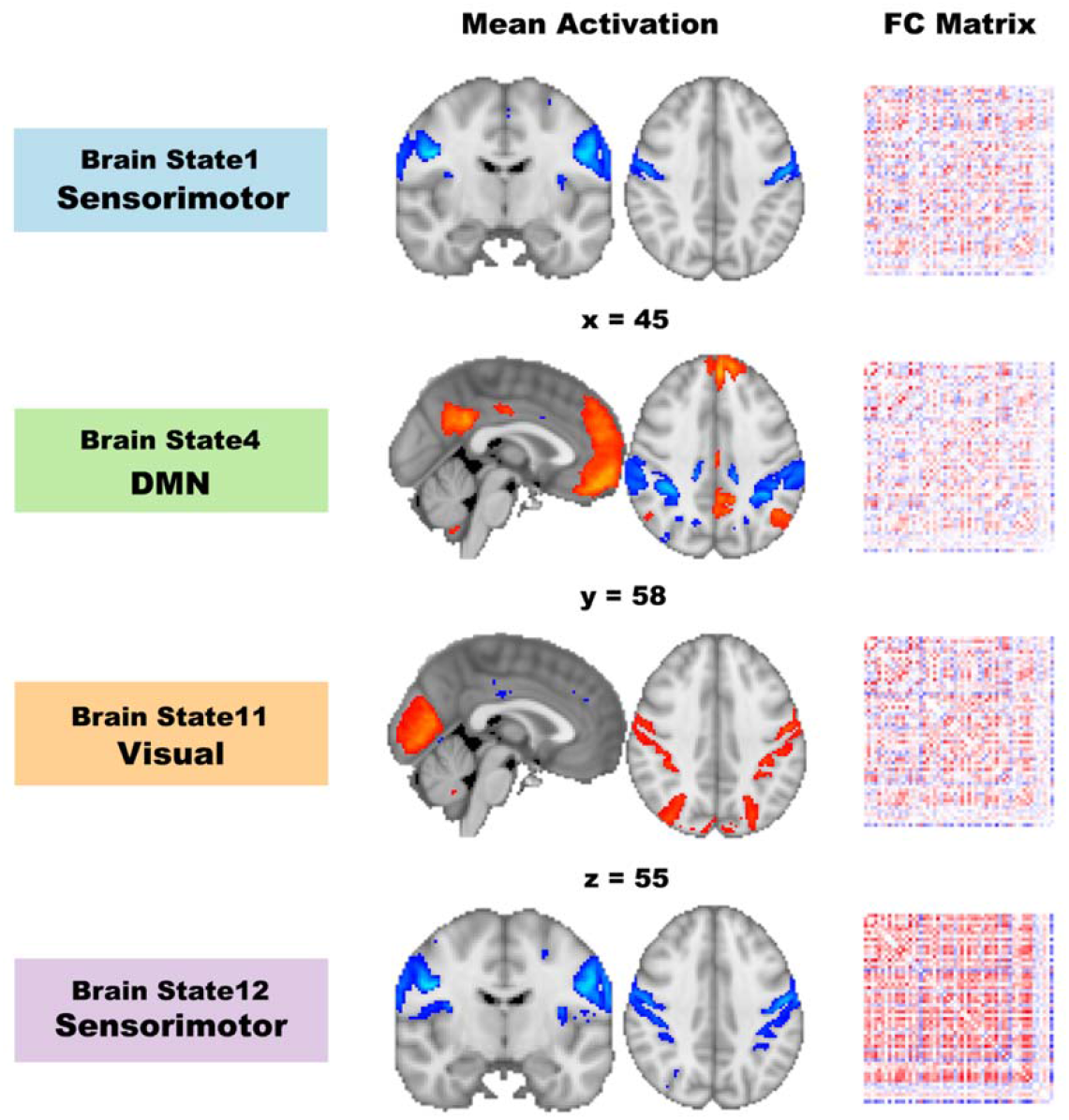
Example Dynamic Brain States Inferred from Hidden Markov Model. A number of brain states common to all participants were estimated with participant-specific time-courses for each individual brain state, which indicate when each brain state is active. These states are characterized by their mean activation and functional matrix. Here, mean activation maps of four example brain states are thresholded to illustrate the top 10% voxels engaged in each brain state, along with their functional connectivity matrices. Network engaged in these brain states are sensorimotor, default mode (DMN) and visual networks, labelled based on the local maximal of connectivity coefficients. Unthresholded mean activation maps for all 12 brain states can be found in Supplemental Material.

**Figure 2.**
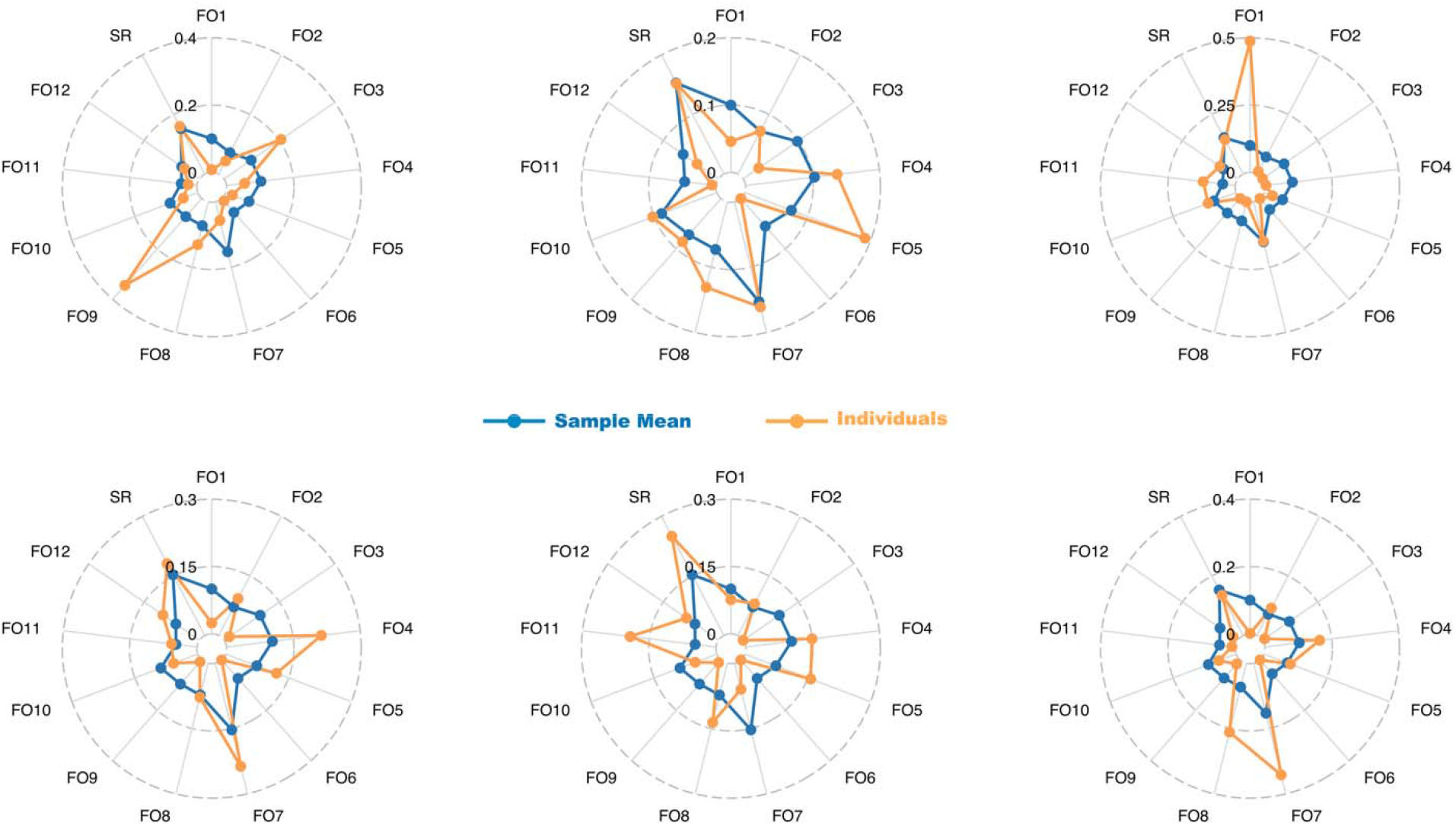
HMM derived brain state measures. Fractional occupancy (FO) per inferred brain state (i.e., indicated by numbers after FO as in FO1) and switching rate (SR) across all brain states were derived for each individual participant to indicate their brain network interactions over time. Data from six randomly selected individual participants were overlaid on the sample mean to illustrate individual variation in these measures.

### Normative range of and abnormality in brain states

Normative modeling with Gaussian process regression was performed to predict each of thirteen brain state measures separately (i.e., FO for each of the 12 brain states and SR across all brain states) from general cognitive ability scores. This approach enabled quantification of individual-specific abnormality scores per brain state measure (i.e., deviations from the estimated normative range). Tests on the group mean of such abnormality scores revealed that the average level of abnormality in FO for seven brain states was significantly different from zero (all p’s<0.0006). Importantly, large individual variation was observed in these abnormality scores (e.g., some participants deviated much more than the others or in different directions), demonstrating unique individual patterns that were not represented by the group mean (*Figure 3*).

**Figure 3.**
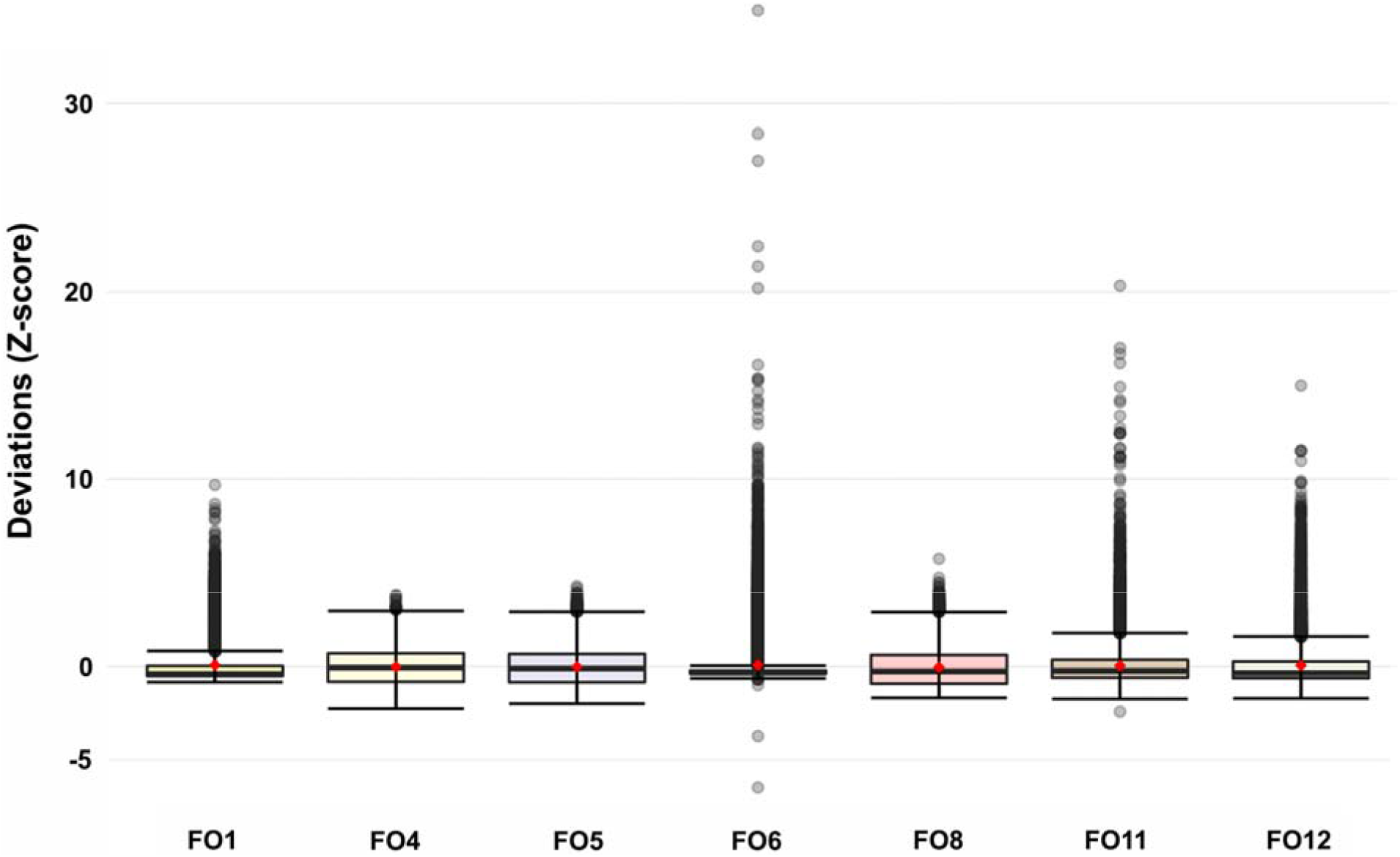
Abnormality in brain states. As shown in the boxplot, averaged deviations (i.e., red dots) in fractional occupancy (FO) for seven brain states were significantly different from zero at the group level with substantial individual variation in both magnitude and direction. Boxes here indicate the inter-quantile range (i.e., middle 50% of the data) and black horizontal lines inside each box represent the median values. Outliers for each FO measure are shown as gray dots beyond the either end of the boxes.

### Individual abnormality in brain states and mental health

To capture the individual level relationship between the cognition-derived brain state measures and mental health, we linked the abnormality scores of each of the seven FOs that were significantly different from zero to the overall mental health scores. Four of these FO abnormality scores showed significant associations with overall mental health after Bonferroni correction for multiple comparisons (t_FO1_=-5.66; t_FO4_=-3.26; t_FO11_=-3.63; t_FO12_=2.93; all p’s<0.0035; *Figure 4*). We further included all four FO abnormality scores in one regression model and the results indicated unique predictive effects of three FO abnormality scores on overall mental health (t_FO1_=-5.01; t_FO4_=-4.31; t_FO11_=-2.02; all p’s<0.009).

**Figure 4.**
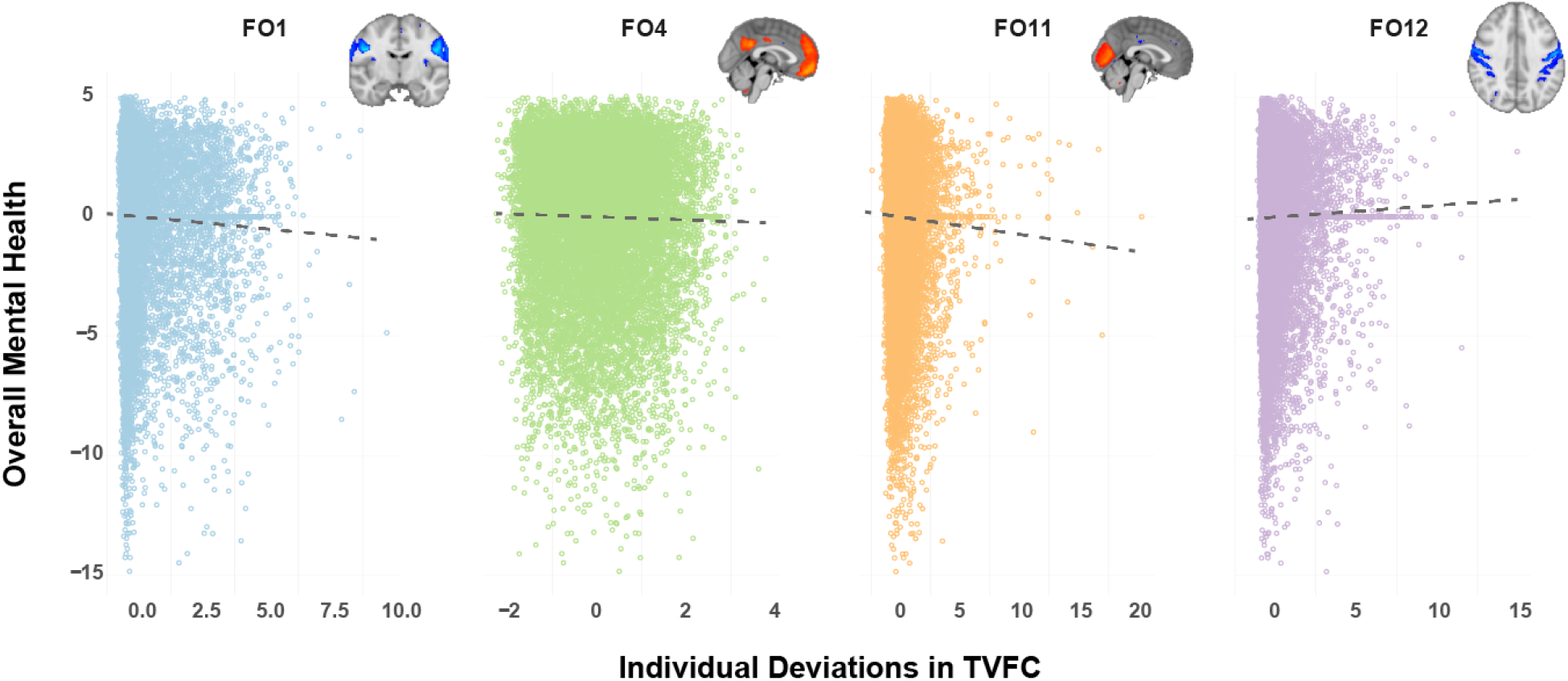
Overall mental health and abnormality in brain states. Individual deviations from the normative range (i.e., abnormality) of fractional occupancy for brain state that engaged sensorimotor (FO1 & FO12), default mode (FO4) and visual networks (FO11) were significantly associated with individual’s overall mental health. Brain images are mean activation maps of each brain state, thresholded to illustrate the top 10% voxels.

As a comparison, we also conducted a more traditional moderator effect analysis to assess whether mental health moderates the relationship between cognitive ability and HMM-derived brain state measures. Importantly, these tests focused on the commonalities across all participants, assuming that interactions between mental health and cognitive ability will impact brain states in the same way for each individual. However, no significant moderator effects were observed (all p’s>0.15).

## Discussion

In this study, we adopted a personalized approach to quantify individual brain state measures that indicate brain network interactions over time, as a function of cognitive ability. Our aim was to test whether individuals’ deviations in brain states from the normative range can be explained by their mental health. Applying normative modeling to a population cohort, we identified several brain states with significant deviations across all participants from the normative range. More importantly, we demonstrated that the degree of abnormality in the fractional occupancy of four brain states could be attributed to the inter-individual difference in overall mental health.

While abnormality in cognitive functions and alterations in the relevant neural correlates have been implicated in a variety of neuropsychiatric conditions, the vast majority of investigations so far has predominantly adopted a case-control approach or divided samples into phenotypic subgroups (e.g., based on symptom scores), which essentially overlooks unique individual characteristics. In the current study, our approach revealed associations between mental health and brain network interaction patterns that were specific to each individual participant’s cognitive ability, whereas more traditional moderator effect tests that focus on commonalities across participants failed to capture such relationships. These findings echo the goal to adopt more personalized approaches in precision psychiatry (*24*) and are well in line with the National Institute of Mental Health’s Research Domain Criteria initiative (RDoC) to take into account individual’s variability rather than diagnostic groups (*25*).

In our findings, significant deviations across all participants were observed in several brain states that involve the default mode and sensorimotor networks (*Figure S2*). Interestingly, whereas the DMN has been consistently associated with various cognitive processes (*26*–*28*) and altered connectivity of this network has also been suggested to underlie multiple mental illness conditions with substantial cognitive deficits (*29*), sensorimotor networks are predominantly involved in motor related processes. However, increasing evidence shows that sensorimotor networks can also contribute to disorder symptoms related to cognitive functions via its connections with other large-scale brain networks such as the DMN (*30*–*32*).

Our findings further revealed associations between the degree of subject-level abnormality in brain state measures (i.e., FO of several brain states) and inter-individual differences in overall mental health. Interestingly, these associations were mostly negative. Since the overall mental health score was derived from the first principal component over mental health questions and it loaded negatively on most of those questions (*Figure S1B*), the observed negative associations between abnormality in FOs and overall mental health imply that the further away individual’s FO deviate from the population norm, the higher the mental health scores they have in general (i.e., more symptoms). These results imply that one’s mental health may impact brain network interactions over time that provide support to cognitive functions and thus point to potential brain mechanisms likely underlying cognitive deficits across a spectrum of major neuropsychiatric disorders. As mental health questions included in the UKB dataset primarily assess mood disorders and neuroticism (*33*), our findings here can be particularly relevant for understanding the neural mechanisms for these disorders.

Whereas recent neuroimaging studies have mainly applied normative modeling to clinical cohorts and studied extreme deviations in estimated brain traits in relation to symptom levels (*20, 34, 35*), our study extended the application of normative modeling to a large population cohort and demonstrated a continuum in the relationship between individual abnormality in brain traits and their overall mental health state. Our findings align well with prior evidence that cognition can be compromised as a function of mental health and highlights potential mechanisms in the brain.

Our findings should be interpreted in light of several limitations. First, the sample of the current study has a specific lifespan (i.e., between 45-81) and is heavily biased towards a specific ethnicity (i.e., White), which may limit the generalization of our results to other populations with different characteristics. Nevertheless, with careful controls for the potential confounding factors including age, sex, education and income levels, as well as head motion during the fMRI data acquisition, our results provide evidence that dynamic interplays of large-scale brain networks may contribute to cognitive deficits due to mental health. Second, HMM was utilized in the current study to infer dynamic brain states that reoccur over time. It is important to note that HMM-derived measures appear to contain information of both time varying and static functional connectivity, and that our implementation of HMM (i.e., model estimation based on the mean and covariance) may capture additional network characteristics (i.e., node amplitude) (*11*). Although it is beyond the scope of the current study to tease apart the unique contributions of these network properties, previous studies did suggest that temporal dynamics of large brain networks are potentially more sensitive to explain inter-individual differences in cognitive traits (*9*–*11*). Lastly, in our study head motion showed correlations with all observed abnormalities in brain state measures that were significant at the group level (all p’s<0.001), as well as with overall mental health (p=0.018). This may appear concerning as head motion is typically considered and modeled as artefacts particularly in the resting-state fMRI studies. However, increasing studies have observed its association with typical functional connectivity networks in the brain (*36, 37*), demonstrated different patterns of head motion for patients versus healthy control groups or in different age groups (*38*–*40*) and suggested that head motion might have genetic roots (*41, 42*). These findings imply that head motion may contain extra information beyond mere noise that is relevant for our interpretations of imaging studies and that estimating the tendency of head motion in relation to the variables of interest may offer more insight into individual level brain signatures. Nevertheless, the reported findings here are carefully controlled for head motion effects in two-folds: ICA-FIX was performed at the individual subject level during preprocessing, and the averaged framewise displacement was further modeled as a between-subject covariate in all statistical analyses. Furthermore, we assessed the robustness of the reported associations between abnormality in brain state measures and individual’s overall mental health by estimating the potential impact of multicollinearity among predictors (i.e., mental health and motion). Our results indicate low levels of collinearity between our predictor of interest (i.e., mental health) and head motion in their predictive effects on the brain state abnormalities (see details in Supplemental Results). Hence, the relationship between brain state abnormality and overall mental health was significant while accounting for head motion effects.

In conclusion, we show in the current study that individualized brain state measures for cognitive ability can be quantified and that the degree of abnormality in such brain states can be attributed to individual differences in overall mental health. These results shed light on the potential brain mechanisms underlying cognitive deficits in mental illness conditions, which may be investigated in patient cohorts in the future.

## Materials and Methods

### 1. Participants

The UK Biobank is a population cohort dataset with extensive behavioral and demographic assessments, lifestyle, and cognitive measures, as well as high quality imaging data. It recruited adults from the general population across Great Britain between 2006 and 2010 (https://www.ukbiobank.ac.uk). This openly accessible population dataset also aims to obtain multimodal brain imaging for 100,000 individuals before 2023 (*43*). We used data from “instance 2” when the initial neuroimaging data were acquired. Participants were included in the analysis if they had resting-state fMRI data and had sufficient data of self-reported mental health questionnaires and of cognitive tasks to derive scores for overall mental health and general cognitive ability (see below). Consequently, data from N=18,634 were analyzed (M_age_=63.65; N_female_=9,913; see sample demographics in Supplemental Materials). All participants provided informed consent. UK Biobank has ethical approval from the North West Multi-Centre Research Ethics Committee (MREC). Data access was obtained under UK Biobank application ID 47267.

### 2. Data acquisition and processing

#### 2.1 General Cognitive Ability

Several well-established cognitive tasks were performed when UK Biobank participants visited the assessment center to measure their cognitive functions including IQ, verbal declarative memory, executive function, and non-verbal reasoning. All tests were administered on a touch screen computer on the day of scanning. Following the practice from previous studies, PCA was used to extract the major component across multiple cognitive tasks that are representative of general cognitive ability (*44, 45*). The summary outcome measures (e.g., mean reaction time, number of correct puzzles) of each task were entered into the PCA with exclusion of task measures wherein data were missing from a substantial number of participants. In case of categorical items, the proportion of each category or level was calculated. This resulted in a final list of 16 items that were entered into the PCA (see full list of items in *Figure S1*). Prior to PCA, imputation was performed to address missing variables using a regularized iterative PCA algorithm, which involves 1) substituting missing values with initial values such as the mean of the variables with non-missing entries, 2) performing a PCA on the imputed data, 3) using the parameters obtained from PCA to predict a new value for the missing one and 4) repeating steps 2 and 3 until convergence. Importantly, only the first N dimensions of PCA in step 2 are used for subsequent estimations, and parameter N is tuned by cross validation that minimizes the mean square error of prediction (*46*). In the current study, the full range of PCA dimensions was considered to tune the optimal number of N. Additional sensitivity analyses were performed to ensure the reliability of the imputation procedure (see Supplementary Materials section S2). Just like the g-factor that indicates general intelligence, the first PCA component was derived as a proxy measure for general cognitive ability and individual scores were used to indicate general cognition levels per participant. To obtain stable estimates of the non-imaging variables of this study (i.e., scores for general cognitive ability and mental health), the imputation and PCA procedures were implemented using the full sample before partitioning data for normative modeling.

#### 2.2 General Mental Health

On the same day when neuroimaging and cognitive task data were acquired, participants also completed a set of self-report mental health questions on a touch screen. The questionnaire items (40 in total) touched upon lifetime mental health with questions about affective disorders, neuroticism, subjective well-being indicated by happiness, and satisfaction levels with health, family relationships, friends and financial situation (*Figure S1*). Individual variables from the mental health questionnaire were excluded with more than 55% missing values and categorical variables were transformed into percentages, resulting in a final list of 33 items that were entered into a PCA after the same imputation procedure as described above (see full item list in *Figure S1*). For the purpose of inferring an overall state of mental health, we used the individual scores for the first PCA component to approximate the *p*-factor that has been proposed to indicate generalized dimensional symptom levels across psychiatric disorders (*47, 48*).

#### 2.3 Imaging data

Resting-state fMRI was obtained using a multiband sequence with an acceleration factor of 8 (TR=0.735; voxel size=2.4×2.4×2.4mm^3^). Preprocessing for the resting-state fMRI data included motion correction, grand-mean intensity normalization, high-pass temporal filtering, unwarping and denoising (*49*). Full details for imaging acquisition and preprocessing steps can be found in UK Biobank Brain Imaging Documentation (https://biobank.ctsu.ox.ac.uk/crystal/crystal/docs/brain_mri.pdf).

Researchers involved in UK Biobank have centrally developed and shared the imaging-derived phenotypes via the UK Biobank showcase (*50*), one of which is subject-specific time-series of 100 resting-state nodes. These time-courses were derived from a dual regression analysis using a group-level independent component analyses (ICA) map as the input. Out of 100 dimensions, 55 were classified as non-noise functional network works, resulting in a matrix size of 490 × 55 for each participant’s time-series, where 490 is the number of acquired volumes. These data were used in the current study to estimate time-varying functional connectivity.

### 3. Statistical Analyses

#### 3.1 Dynamic Brain States

We derived brain state measures by fitting a Hidden Markov Model (HMM) to the aforementioned fMRI time-series data. In brief, HMM assumes that brain activity can be characterized as a number of discrete brain states. Under this framework, the brain activity at each time point can be modeled as a mixture of Gaussian distributions with each distribution corresponding to a different state (*10, 51*). Importantly, each of the estimated brain states is described by a distinct timeseries and functional connectivity pattern (*52*). HMM can be viewed as a probabilistic description of the data in which the hidden information (i.e., brain state) is captured from observable measures (i.e., fMRI time courses). An increasing number of studies have applied HMM to neuroimaging data, demonstrating it as a promising approach for understanding the temporal dynamics of brain functions (*52*–*55*).

In accordance with previous work applying HMM to fMRI data, we inferred 12 brain states using the resting-state fMRI time-series data from all participants (*52, 56*). Although different numbers can be assumed and hence enforced on HMM for inferring brain states, there is no single “best” state dimensionality as assumptions can vary depending on the specific interest in the level of the brain’s hierarchical organization. To align our study with previous HMM work on the fMRI data, we followed the choice of twelve brain states (52, 56). Nevertheless, the HMM model was repeated multiple times to ensure the stability of the state estimates (see Supplementary Material section S3). Measures including fractional occupancy (FO) that represents the average state probability across time for each brain state, and the switching rate among brain states that indicates the brain state stability of each participant were quantified at the individual level. These measures were calculated based on the inferred probability of brain state occurrence at specific time points and together can be considered as indications of subject-specific patterns of brain states (i.e., a total of 13 measures from the estimated model with 12 brain states).

#### 3.2 Normative modeling of brain state measures as a function of cognition

To obtain statistical inferences at the individual level with respect to the normative range of brain state measures, we used a Gaussian Process Regression (GPR) to predict HMM-derived measures from the general cognitive ability scores. GPR is a non-parametric probabilistic Bayesian method that can provide reliable estimates of uncertainty for the prediction and has been widely applied in the machine learning field (*57*). Using GPR, we estimated each individual measure of brain states separately as a function of general cognitive score, and quantified subject-level deviations from the estimated normative range of per measure (i.e., abnormality score) for each individual participant. No other nuisance variables were considered at this stage. Abnormality scores were calculated as the difference between estimated brain state measure value from the normative model and the actual value derived from HMM, which was further divided by the estimated standard deviation from the normative model (i.e., Z scores) (*20*).

To infer the normative range with the current sample size (N>18K), we leveraged local approximate GP modeling. In contrast to full GPR that requires obtaining a full set of predictive equations, local approximate GPR involves approximating the predictive equations at a particular location (e.g., one individual cognitive ability score in our case) via a subset of the data. As the remote elements from the subset to th location have vanishingly small influence on prediction, ignoring these elements in order to work with much smaller matrices can be computationally efficient with little impact on prediction accuracy (*58*). For a more detailed description about localized approximate GPR, please see the Supplementary Material, section S4. 10-fold cross validation was employed to ensure generalizability of the estimates, where data were partitioned into training and testing sets across participants per fold (i.e., one out of the total ten subsamples was held for testing per fold).

Once the abnormality scores for all participants were estimated, one-sample t-tests were used to examine which of the brain state measures had abnormality scores that on average were significantly different from zero, using the adjusted P-value to account for multiple comparisons (i.e., p<0.05/13=0.00385). Only the brain state measures for which the group mean abnormality was significantly different from zero were included in subsequent analyses, linking brain state abnormality to mental health.

#### 3.3 Linking brain state abnormality to mental health

In order to examine whether inter-individual differences in the degree of abnormality in brain state measures can be attributed to overall mental health, we performed linear regression models. Abnormality scores were included as the independent variable and the overall mental health scores the dependent variable. We further considered age, sex, education and income levels, as well as head motion during resting-state imaging acquisition (i.e., the mean framewise displacement) as nuisance covariates and included those in the analyses that showed significant correlations with overall mental health scores. Multiple comparisons were accounted for the Beta values of abnormality scores (i.e., predictor) in each model using Bonferroni correction (i.e., adjusted p<0.05/7=0.007). To examine whether any abnormality scores of brain state measures exhibited unique association with mental health over and above each other, we further conducted a multiple regression model including all brain state abnormality scores that passed the one-sample t-test.

#### 3.4 Implementation of statistical analyses

All statistical analyses were conducted using R version 3.6.1 (R Core Team, 2019). Imputation prior to PCA was conducted for missing values using the R function imputePCA from missMDA package (*46*), and function prcomp from basic stats package (*59*) was used for the PCA analyses. Local approximate GPR was performed using the laGP package (*60*).

#### 3.5 Data and materials availability

The UK Biobank data that support the findings of this study can be accessed by researchers on application (https://www.ukbiobank.ac.uk/register-apply/). Variables derived specifically for this study will be returned to the UK Biobank for future applicants to request. R scripts for normative modeling can be found on GitHub (https://github.com/PersonomicsLab/laGP_nm). Matlab scripts for running HMM are also available on GitHub (https://github.com/OHBA-analysis/HMM-MAR).

## Acknowledgments

We are grateful to UK Biobank and the UK Biobank participants for making the resource data possible, and to the data processing team at Oxford University for producing the shared processed data. JDB is funded by the NIH (1 R34 NS118618-01) and the McDonnell Center for Systems Neuroscience.

